# Role of astrogial Connexin 43 in pneumococcal meningitis and pneumolysin cytotoxicity

**DOI:** 10.1101/2020.01.15.907923

**Authors:** Chakir Bello, Yasmine Smail, Vincent Sainte-Rose, Isabelle Podglajen, Alice Gilbert, Vanessa Oliveira Moreira, Fabrice Chrétien, Martine Cohen Salmon, Guy Tran Van Nhieu

**Affiliations:** Team Intercellular Communication and Microbial Infections, Center for Interdisciplinary Research in Biology, Collège de France, 75005 Paris, France; Institut National de la Santé et de la Recherche Médicale U1050, 75005 Paris, France; Centre National de la Recherche Scientifique UMR7241, 75005 Paris, France; MEMOLIFE Laboratory of excellence and Paris Science Lettre; Team Physiology and physiopathology of the gliovascular unit, Center for Interdisciplinary Research in Biology, Collège de France, 75005 Paris, France; Experimental Neuropathology Unit, Institut Pasteur, 75015, Paris, France

## Abstract

*Streptococcus pneumoniae* or pneumococcus (PN) is a major causative agent of bacterial meningitis with high mortality in young infants and elderly people. The mechanism underlying PN crossing of the blood brain barrier (BBB) remains poorly understood. Here, we show that the gap junctional component connexin 43 expressed in astrocytes (aCx43) plays a major role in PN meningitis. Following intravenous PN challenge, mice deficient for aCx43 developed milder symptoms and showed severely reduced bacterial counts in the brain. We show a role for aCx43 in the PN-induced fragmentation of astrocytic GFAP filaments associated with bacterial translocation across endothelial vessels and replication in the brain cortex. aCx43 triggers the PN- and Ply-dependent GFAP fragmentation and nuclear shrinkage in *in vitro* cultured astrocytes. We showed that purified pneumolysin (Ply) co-opted Cx43 to promote the permeabilization and cytosolic calcium (Ca^2+^) increase of host cells, a process sensitive to extracellular ATP depletion. These results point to aCx43 as a major player during bacterial meningitis and extend cytolytic mechanisms implicating other host cell plasma membrane channels proposed for small pore-forming toxins, to Ply, a cholesterol-dependent cytolysin, at concentrations relevant to bacterial infection.

---

Numerous PN meningitis-associated factors have been reported, that target host cell receptors, degrade the extracellular matrix or intercellular junctional components, trigger inflammatory responses and allow escape from innate defence mechanisms [1-3]. PN may cross the BBB via a paracellular route, following the loosening of endothelial cell junctions triggered by the pore-forming toxin (PFT) Pneumolysin (Ply) and inflammatory responses[2]. PN can also transcytose through brain vascular endothelial cells by targeting receptors such as the Platelet-Activating Factor receptor (PAFR), the poly-Immunoglobulin Receptor (pIgR) or the Platelet Endothelial Cell Adhesion Molecule (PECAM-1)[3, 4]. Interactions between PN and these endothelial cell receptors require down-regulation of the bacterial capsule, essential for bacterial survival in the blood and evasion of phagocytosis [3].

Ply is a critical virulence factor involved in all PN invasive diseases including meningitis [3, 5]. Ply may not be required in models of PN meningitis following direct intra-cranial injection of bacteria, but *ply* mutant do not promote meningitis following intravenous injection suggesting a role in PN crossing of the BBB. Ply may favor PN translocation by triggering inflammation that fragilizes the BBB [6]. Ply was shown to alter astrocyte shape, glutamate signaling and to trigger synaptic damage at non-lytic concentrations and during PN meningitis [7-9], however, its role during the early phase of PN translocation across the BBB remains ill defined.

The large majority of cellular studies on PN translocation across the BBB have been performed using tissue culture brain endothelial cells. While brain endothelial cells represent the mechanical barrier isolating the brain cortex from the bloodstream, other cell types forming the neurovascular unit regulate the BBB function [10, 11]. Among these, astrocytes form glial networks in the various brain compartments and extend end-feet contacting vessels of the brain vasculature [12, 13]. The astrocytes at this interface modulate the integrity and functions of the blood-brain barrier, neuroinflammation, cerebral blood flow, and interstitial fluid drainage [13-15].

Using mice retro-orbital vein injection, we found that all aCx43^FL/FL^ control mice developed meningitis when challenged with TIGR4, a PN serotype 4 wild-type strain [16]. All PN infected mice showed reduced activity, associated with piloerection at 9 H post-infection, with aggravating symptoms including hunched postures and absence of motility at 24 H post-infection, the longest incubation time at which mice were sacrificed to limit animal suffering (Figs. 1A and S1). As shown in Fig. 1B, CFU determination in brains of aCx43^FL/FL^ mice indicated that bacterial translocation was detected as early as 3 H post-injection and increased exponentially over the 24 H incubation period to reach a median value of 6.2 × 10^4^ CFUs / mg, a time point at which rupture of the BBB integrity could be detected macroscopically (Fig. S1A). Bacterial CFUs could be detected in the cerebrospinal fluid at 24H but not at 3H post-infection, suggesting the brain as a primary infection target (Fig. S1B). In control experiments, aCx43^FL/FL^ mice showed no symptoms when infected with irrelevant an *E. coli* K12 strain and no CFU counts could be detected in sampled brains at 24 H post-infection (N = 4). Interestingly, aCx43^-/-^ mice did not present obvious symptoms or only slightly reduced activity up to 24 H post infection, suggesting that infection was controlled. Accordingly, PN translocation in the brain of aCx43^-/-^ mice was significantly decreased with a median value of 1.2 × 10^2^ CFUs / mg at 24 H post-infection. Blood counts shared similar median values between aCx43^FL/FL^ and aCx43^-/-^ mice during the first 13H incubation. However, at 24 H post-infection, aCx43^FL/FL^ mice showed blood titers that were 700-times higher than aCx43^-/-^ mice (Fig. 1C), consistent with secondary systemic infection from the brain. PN infection led to up-regulation of the pro-inflammatory cytokines TNF-α and IL1-β, as well as of endothelial inflammation markers as revealed by qRT-PCR, but to similar extent in aCx43^FL/FL^ and aCx43^-/-^ mice despite the difference in bacterial titers (Fig. S2).

**Fig. 1.**
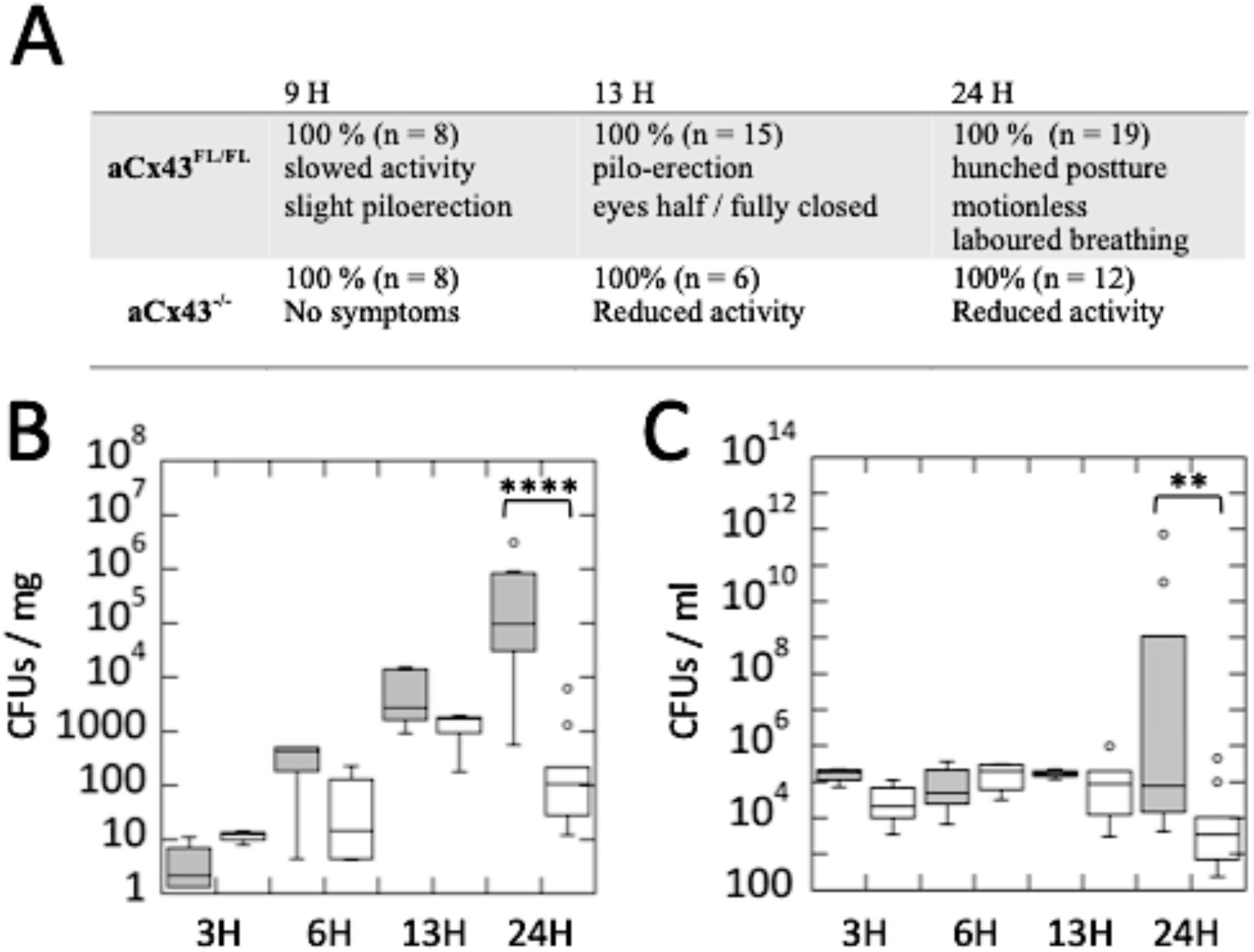
aCx43 involvement in PN meningitis. **A**, symptoms of C57/BL6 mice proficient (aCx43^FL/FL^) or deficient (aCX43^-/-^) for astroglial Cx43 following intravenous PN challenge. **B, C**, box plots showing medians of CFU determination associated with brain (B) or blood (C) samples at the indicated post-infection time. Grey boxes: aCx43^FL/FL^ mice; empty boxes: aCX43^-/-^mice. **B**, aCx43^FL/FL^: 3H, N = 4; 6H, N = 4, 13H, N = 5; 24H, N = 12. aCx43^-/^: 3H, N = 4; 6H, N = 4, 13H, N = 3; 24H, N = 12. **C**, aCx43^FL/FL^: 3H, N = 4; 6H, N = 4, 13H, N = 5; 24H, N = 9. aCx43^-/-^ 3H, N = 4; 6H, N = 4, 13H, N = 3; 24H, N = 9. Wilcoxon test. **: p = 0.01; ****: p < 0.0001.

To further characterize the early events associated with PN translocation across the BBB, we performed brain immunofluorescence analysis (Methods). No bacteria were detected in the brain cortex at 3 H post infection and only 0.7 to 3.3 bacteria or bacterial clusters / mm^2^ were observed at 6 H and 13 H post-infection. As shown in Fig. 2A, bacteria were found in close association with brain vessels, consistent with early brain translocation events. Immuno-labeling revealed the presence of small capsular remnants at the vicinity of bacteria associated with brain vessels (Fig. 2A). Capsular remnants associated with bacteria were detected in the lumen of brain vessels, suggesting that bacterial lysis or capsular shedding reported during *in vitro* interaction with endothelial cells also occurred in the brain vasculature (Fig. 2A, arrowhead; [2, 17]). Remarkably, such capsular remnants were also detected in the brain cortex seemingly leaking from vessels in association with translocated bacteria at 6H post-infection, consistent with loss of vessel integrity at early stages of crossing of the BBB by PN (Fig. 2A, arrows). Capsular remnants and loss of endothelial vessel integrity was clearly detected in association with bacterial clusters at 13H post-infection. Consistent with the scoring of CFUs from sampled brains, aCx43^-/-^ mice showed three times less translocation events compared to aCx43^FL/FL^ mice as early as 6H post-infection (Figs. 2B, C). Single or a discrete number of bacteria were observed, with lesser capsular remnants and loss of vessel destruction, consistent with poor PN translocation across the BBB in aCx43^-/-^ mice (Fig. 2B). Bacterial cluster size quantification also indicated that PN brain intra-cortical replication was higher in aCx43^FL/FL^ relative to aCx43^-/-^ mice, with clusters 6 and 23-times bigger at 6H and 13 H post-infection, respectively (Figs. 2C, D).

**Fig. 2.**
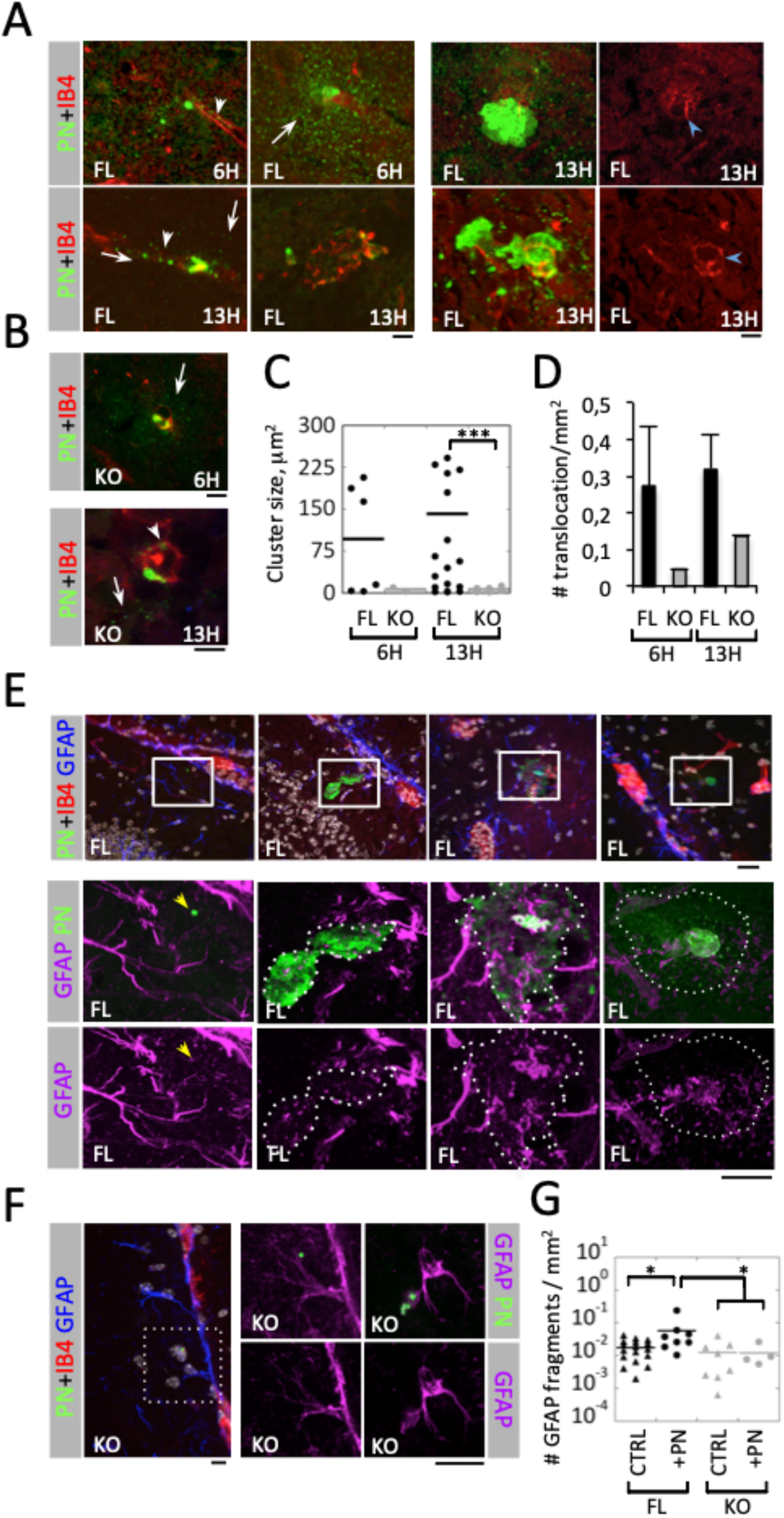
aCx43 potentiates PN translocation across the BBB and fragmentation of astrocytic GFAP processes. Immunofluorescence analysis of brain slices from PN-infected mice. The time post-infection is indicated. Scale bar = 5 μm. **A, B, E, F**, red: IB4-labeling of endothelial vessels; green: PN capsule. FL: aCx43^FL/FL^ mice; KO: aCx43^-/-^ mice. **A, B**, capsular remnants in the brain cortex (white arrows) or in vessels (white arrowheads). **A**, right panels: blue arrowheads point at vessel damages. **C**, size of translocated PN microcolony. 6H: FL, N = 3, 6 foci; KO, N = 3, 2 foci. 13H: FL, N = 3, 20 foci; KO, N = 3, 9 foci). **D**, frequency of bacterial translocation events per mm^2^ of brain slice (N = 2, n = 900 60x microscopy fields). **C, D**, median values are indicated. **E**, top panels: blue: GFAP staining; bottom panels: higher magnification of the inset shown in the top panel. Purple: GFAP staining. The yellow arrow points to the GFAP association with a single translocated bacterium. **F**, quantification of GFAP fragments per μm2 in area corresponding to capsular shedding (dotted area) associated with translocated bacterial microcolony. FL CTRL, N = 2, > 10 000 fragments; FL+PN, N = 2, 1658 fragments; KO CTRL, N = 2, 3619 fragments; KO+PN, N = 2, 263 fragments. Mann and Whitney. *: p < 0.05.

Strikingly, bacterial intracortical growth in aCx43^FL/FL^ mice brains was associated with the fragmentation and GFAP astrocytic filaments (Fig. 2E, dotted area). In association with the disruption of the GFAP network, nuclear shrinkage and fragmentation could also be detected in astrocytes associated with intracortical PN microcolonies reminiscent of the action of Ply (Fig. S3) [18]. The fragmentation of GFAP filaments not only occurred in close contact to bacterial microcolonies but also at the vicinity defined by the area corresponding to diffusion of shed capsular materials (SCMs) (Figs. 2E and S3, dotted area) suggesting that an action of secreted bacterial products. Such fragmentation was seldom observed in aCx43^-/-^ mice brains, where single translocated bacteria were detected in close association with GFAP labeled astrocytic processes (Fig. 2F). Consistently, quantification based on the size and shape of GFAP-labeled structures indicated that fragmentation of GFAP-labeled filaments was more pronounced in the area defined by SCMs compared to fields devoid of bacteria in aCx43^FL/FL^ mice brains (Figs. 2G, S3). In contrast, no difference in GFAP structures fragmentation was observed in astrocytes associated with translocated bacteria in aCx43^-/-^ mice brains (Figs. 2G, S3).

aCx43 is a main Cx expressed in astrocytes forming hemichannels involved in paracrine signaling particularly relevant for the regulation of the BBB permeability [19, 20]. Our results indicated a role for aCx43 in the BBB translocation by PN in association with a loss of brain vascular endothelial integrity and astrocytic damages.

To further characterize the role of aCx43 during PN infection, mice cortical primary astrocytes were isolated and challenged *in vitro* with PN (Methods). As shown in Figs. 3A and S4, following PN challenge for 90 min, a decrease in the number and length of GFAP processes was detected in aCx43^FL/FL^ and aCx43^-/-^ astrocytes compared to non-infected cells (Fig. S4). As observed in brain slices, a clear fragmentation of the GFAP network was detected in cultured aCx43^FL/FL^ astrocytes (Figs. 3A, B), that was not detected in control unchallenged aCx43^FL/FL^ astrocytes and PN-challenged aCx43^-/-^ astrocytes (Figs. 3A, B), indicating a role for aCx43. In instances, GFAP fragmentation in aCx43^FL/FL^ astrocytes was associated with nuclear shrinking (Fig. 3A), a process that was more pronounced following bacterial challenge for 4 hours (Fig. 3C, arrows). At this time point, nuclear shrinkage in aCx43^FL/FL^ astrocytes was associated with cell retraction and (Figs. 3A, B), as reported for the cytotoxic action of the pore-forming activity of Ply on neuronal and microglial cells [8, 18]. Consistently, when used as a proxy, no detectable nuclear shrinkage was observed for a Ply-deficient isogenic PN mutant (Figs. 3C, D). Strikingly, nuclear shrinkage was less obvious in astrocytes derived from aCx43^-/-^ mice upon challenge with wild-type PN or aCx43^FL/FL^ astrocytes in the presence of the Cx channel inhibitor carbenoxolone (Figs. 3C, D), suggesting a role for aCx43 in Ply-mediated cytotoxicity. Consistently, nuclear shrinkage was phenocopied when cells were challenge with purified recombinant Ply at concentrations up to 500 nM (Fig. 3D), an effect that was inhibited by carbenoxolone (Fig. 3E).

**Fig. 3.**
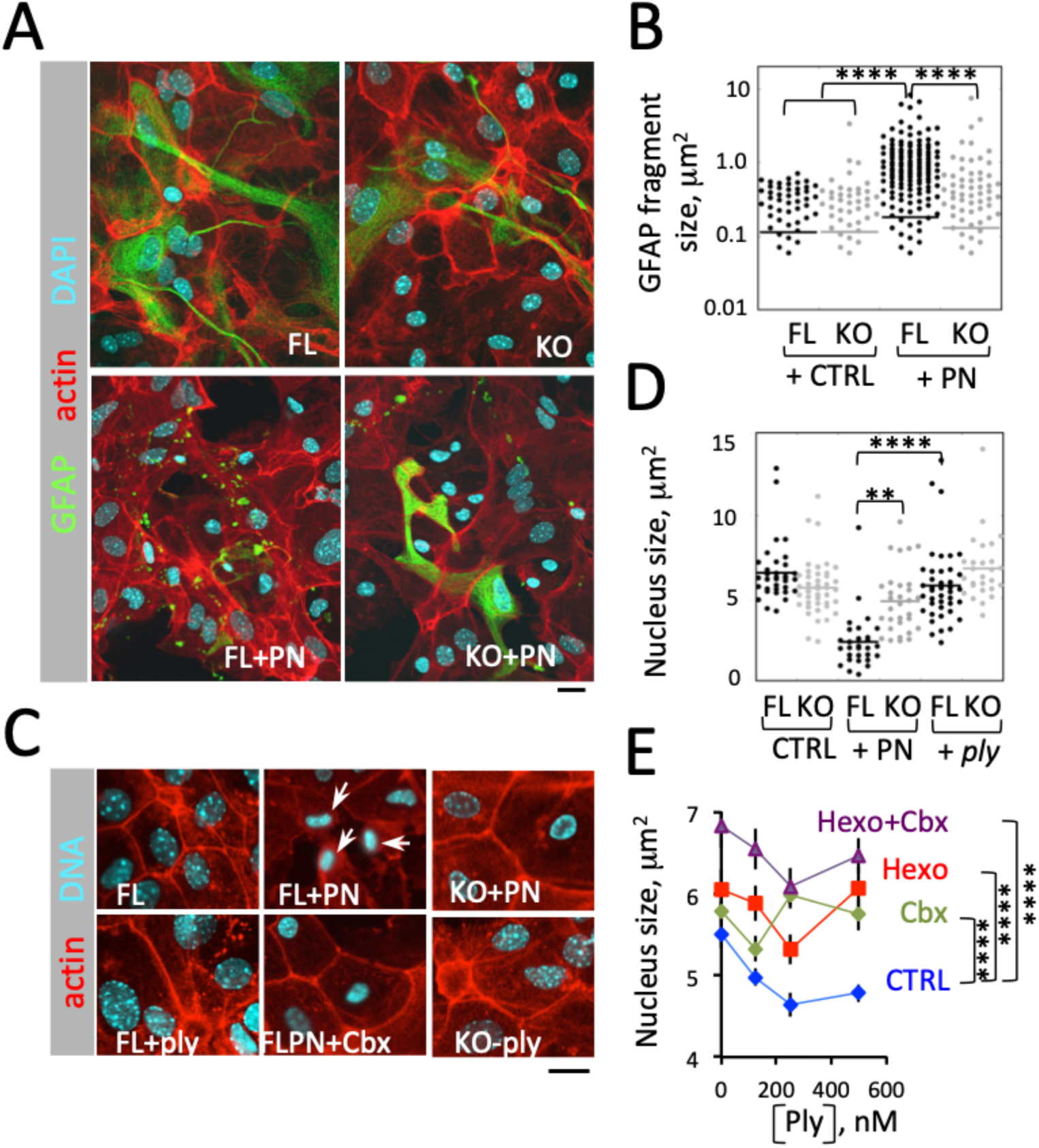
Ply mediates aCx43-dependent GFAP fragmentation and nuclear shrinkage in astrocytes. *in vitro* cultured astrocytes derived from brains of wild-type (FL) or aCx43^-/-^ (KO) mice were challenged with wild-type TIGR4 (PN) or an isogenic *ply* mutant (*ply*) (**A-D**), or purified Ply at the indicated concentrations for 60 min (**E**). Samples were fixed and processed for immunofluorescence staining of GFAP (green), F-actin (red) or nuclear DNA using DAPI (cyan). Cbx: challenge in the presence of 100 μM carbenoxolone. Scale bar = 5 μm. PN challenge for: **A, B**: 90 min; **C, D**: 4 hours. **B**, quantification of GFAP fragmentation. The bars represent median values. FL: N = 2, 78 cells, 9196 fragments; KO: N = 2, 93 cells, 2088 fragments; FL+PN: N = 2, 131 cells, 3584 fragments; KO+PN: N = 2, 103 cells, 1827 fragments. **C**, Arrows point at nuclear shrinkage. **D, E**, representative experiments of nuclear size quantification. Bars: median values. Mann and Whitney. **: p < 0.01; ****: p < 0.0001. **E**, + Hexo: incubation in the presence of hexokinase. ANOVA ****: p < 0.001.

Ply-mediated cytotoxicity has been linked to mitochondrial dysfunction and cytosolic Ca^2+^ increase [18, 21]. Our results suggested that Cx43 channels play a major role in Ply-mediated cytotoxicity. These findings are reminiscent of cytotoxicity induced by the small pore forming toxins RTXs involving the release of ATP through plasma membrane channels formed by purinergic P2X7 receptors and pannexins [22][23]. To further characterize the mode of action of Ply and extend findings to cells other than astrocytes, we analyzed the effects of Ply in HeLa cells that do not express known Cxs and HeLa cells stably transfected with Cx43 (HCx43; [24]). As shown in Figs. 4A, B, challenge with PN did not show to detectable change in morphology of parental HeLa cells, but cell retraction was clearly observed for HCx43 cells. As expected, cell retraction was dependent on Ply, since it was not detected upon challenge with *ply* mutant and was induced by purified Ply (Figs. 4A-C). As observed for small PFTs, Ply-induced rounding of HCx43 cells was inhibited by hexokinase that depletes extracellular ATP (Fig. 4D). Consistent with increased plasma membrane permeabilization linked to ATP release through Cx43 hemichannels, in dye release assays, the rates of calcein fluorescence decrease were significantly higher in HCx43 cells compared to parental HeLa cells or cells challenged in the presence of hexokinase (Figs. 4E and S5). Low concentrations of Ply triggered Ca^2+^ responses in HCx43 cells that were not observed in HeLa cells (Figs. 4F and S6). Also, high concentrations of Ply triggered a lasting increase in intracellular Ca^2+^ consistent with Ca^2+^ influx linked to plasma membrane permeabilization three-times more frequently in Hcx43 cells compared to HeLa cells (Fig. S6).

**Fig. 4.**
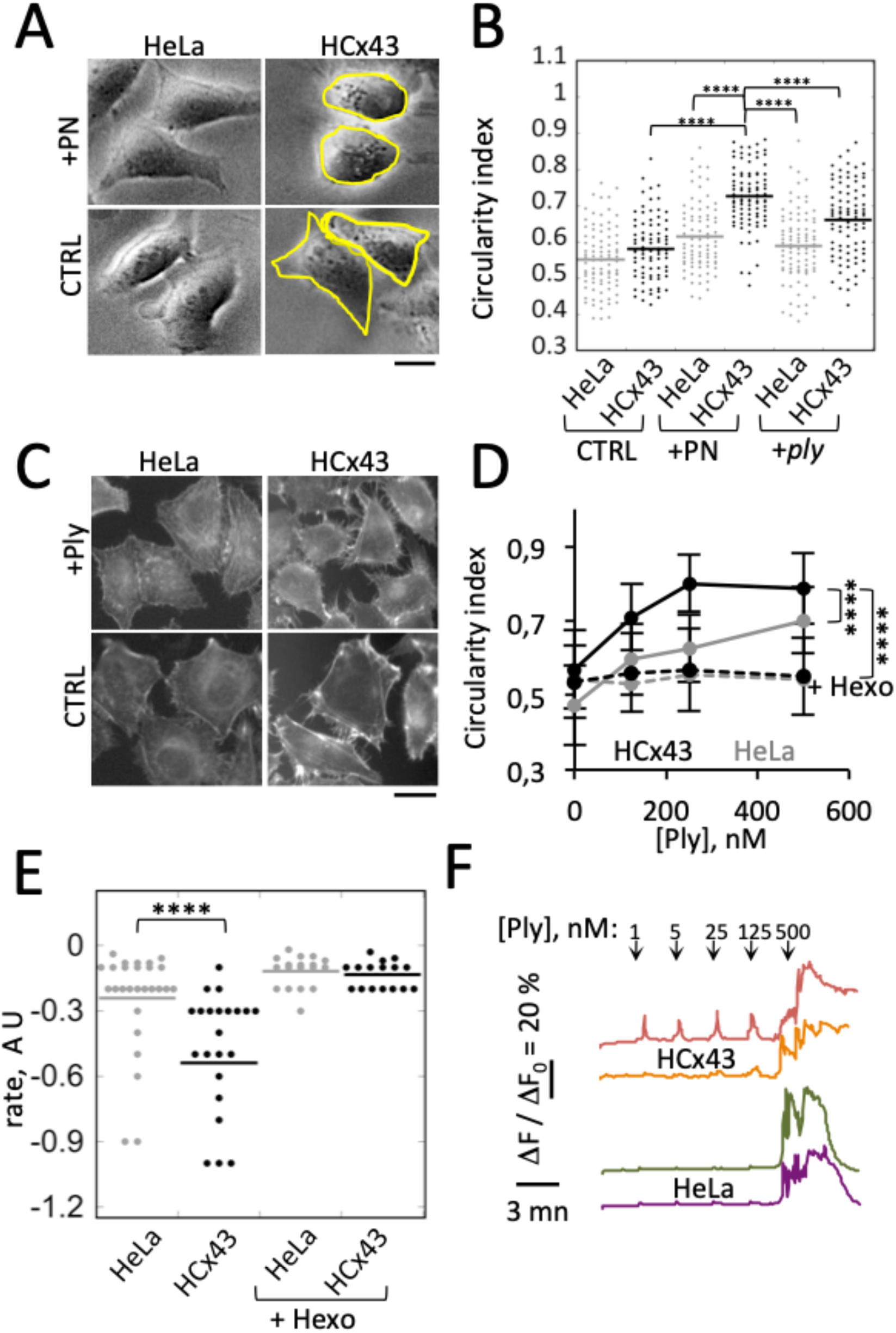
Cx43 expression confers Ply-mediated cytotoxicity. Parental HeLa cells or stable transfectants expressing Cx43 (HCx43) were challenged for 90 min with wild-type TIGR4 (+PN) or an isogenic *ply* mutant (+ *ply*) (**A, B**), or with purified Ply (+Ply) (**C-G**). CTRL: non-infected cells. Scale bar = 5 μm. **A**, representative phase contrast images. Cell contours are drawn in yellow in left panels. **B, D**, representative experiments of the quantification of cell retraction using circularity index as a proxy. Median values are represented. **B**, > 25 cells per sample. Mann-Whitney. ****: p < 0.001. **D**, > 72 cells per sample. ANCOVA. ****: p < 0.001. **C**, samples were fixed and processed for fluorescence staining of F-actin. **D, E**, + Hexo: incubation in the presence of hexokinase. **E**, representative experiment of Ply-induced calcein release. Calcein-loaded cells were incubated with 250 nM Ply. The rates of calcein decrease were determined from linear fits (Methods). Bars: median values. Mann-Whitney. ****: p < 0.001. **F, G**, cells were loaded with the Ca^2+^ indicator Fluo-4. **F**, representative traces of Ca^2+^ variations in single cells. The arrows indicate the addition of Ply at the indicated concentrations.

As a Cholesterol-Dependent Cytolysin (CDC), Ply is believed to cause cytotoxicity through the formation of large pores in host cell plasma membranes [5, 25]. However, our findings support the notion that at physiological concentrations, Ply cytotoxicity involves aCx43 hemichannels and the release of extracellular ATP to promote host cell permeabilization. These findings are consistent with published reports showing that low concentrations of Ply may induce the formation of small pores in host cell membranes, perhaps linked to the formation of incomplete rings or arcs [26, 27]. Thus, Ply may function in a manner similar to the small pore forming toxins RTXs shown to require host cell plasma membrane channels and ATP release to induce hemolysis [28]. The characterization of Cx43 as major actor in Ply-mediated cytotoxicity has major implications for PN meningitis. Based on our findings, we propose that during the early stages of PN meningitis, PN triggers loss of vascular endothelial cell integrity (Fig. S7). This loss of integrity is favored by the targeting of astrocytes by secreted PN products. Among PN secreted products, Ply targets astrocytes in a process dependent on aCx43 hemichannels and eATP release (Fig. S7). Because PN growth is subjected to stringent requirements, leakage from the brain vasculature may also provide with blood nutriments favoring PN intracortical growth in aCx43-proficient but not deficient mice (Fig. S7). Future works is required to appreciate the role of secreted PN products and hemichanel-mediated paracrine signaling at the various stages of bacterial meningitis.

## METHODS

### Bacterial strains, cell lines, and reagents

The *ply* isogenic mutant from the S. pneumonia serotype 4 clinical isolate TIGR4 was a kind gift from Andrew Camilli (Tufts University, Boston, USA). Bacteria were grown in Todd Hewitt Broth containing (#BD249240, Thermofisher) 0.5% Yeast Extract at 37°C (#210929, Thermofisher) and plated on Columbia blood agar plates (# 43041, Biomérieux, France). Primary astrocytes were derived from mice forebrains as previously described (ref). HeLa cells (ATCC CCL-2™) were from ATCC, and the stable HeLa cell line expressing human Cx43 were described previously [24]. Cells were grown in DMEM (Dulbecco’s Modified Eagle Medium, #10567-014, Thermofisher) containing 10 % fetal calf serum in a 37°C incubator supplemented with 10% CO_2_. The anesthetics Imagen (Ketamin) was from Merial and Rompun (Xylazin) was from Bayer Heathcare). The rabbit polyclonal anti-pneumococcal serotype 4 capsular antibody was from Statens Serum Institute, Copenhagen, Denmark). The mouse monoclonal anti-GFAP (Glial Fibrillary Acidic Protein) antibody (# G3893) and Alexa Fluor 563-conjugated isolectin IB4 (# I21412), secondary goat anti-mouse IgG antibody conjugated to Alexa Fluor 488 (#A11029), goat anti-rabbit IgG conjugated to Alexa Fluor 555 (A21424), and Alexa Fluor 633 Phalloidin (#A22284), Fluo4-AM calcium indicator (#F14201), calcein-AM (#C3100MP) were from Thermofisher Scientific. 2-(4-Amidinophenyl)-6-indolecarbamidine dihydrochloride, 4’, 6-Diamidino-2-phenylindole dihydrochloride (DAPI, #D9542), hexokinase (#9001-51-8), carbenoxolone (# C4790) were from Sigma Aldrich.

### Cloning and purification of recombinant Ply

Ply was cloned into the TOPO-TA cloning vector pET101D (#K10101, Thermofisher) using the following primers: 5’-CACCATGGCAAATAAAGCAGTAAATGAC-3’ and 5’-GTCATTTTCTACCTTATCCTCTACCTGAGG-3’. The insert was verified by DNA sequencing. Purification of recombinant Ply was performed using Talon resin (#PT1320-1, Clontech Laboratories Inc.) affinity chromatography from freshly transformed BL21/DE3 *E. coli* following the manufacturer’s instructions. Samples were store in 25 mM HEPES, 50 mM NaCl, 0.1% beta-mercaptoethanol in aliquots at −80°C and defrosted freshly before use.

### Mice meningitis model

Experiments and techniques reported here complied with the ethical rules of the French agency for animal experimentation and with the Institute of Medicaments, Toxicology, Chemistry, and the Environment animal ethics committee (Paris Descartes University, Agreement 86-23). aCx43^-/-^ mice deficient for astrogial Cx43 and respective proficient aCx43^FL/FL^ mice were described previously [29]. PN cultures were freshly grown to OD_600nm_ = 0.2, and resuspended in PBS buffer at a final concentration of 5 ×10^8^ cfu/ml. Following anesthesia, 6-9 weeks old C57BL/6 mice were infected through intravenous retro-orbital injection by 50 uls of the bacterial suspension. At various time points post-infection, mice were anesthetized and blood was sampled for CFU determination. For the macroscopic analysis of the BBB integrity, mice were injected via retro-orbital vein with 60 μl of PBS containing 2% Evans Blue. Mice were subjected to intracardiac perfusion with 20 mls of sterile PBS using a peristaltic pump at a flow rate of 2.5 mls / mn prior to brain sampling. Sampled brains were either flash-frozen at −80°C for subsequent RNA extraction and qRT-PCR analysis, immediately homogeneized for CFU determination following plating on blood agar plates, or processed for cryosection and immunofluorescence analysis.

### QRT-PCR analysis

Total RNAs were isolated from frozen brain samples homogenized in Trizol (Life Technologies) and chloroform using glass beads and the RNeasy Lipid Tissue kit (Qiagen Corp.). qRT-PCR was performed using the Superscript II reverse transcriptase kit (Invitrogen) and SYBR green PCR master kit (Applied Biosystems) and the following pairs of primers for TNF-α: 5’-GACCCTCACACTCAGATCATCTTCT-3’ and 5’-CCTCCACTTGGTGGTTTGCT-3’; IL-1b: 5’-CTGGTGTGTGCAGTTCCCATTA-3’ and 5’-CCGACAGCACGAGGCTTT-3’; I L-1Ra: 5’-CTTTACCTTCATCCGCTCTGAGA-3’ and 5’-TCTAGTGTTGTGCAGAGGAACCA-3’; vimentin: : 5’-CGGAAAGTGGAATCCTTGCA-3’ and 5’-CACATCGATCTGGACATGCTGT-3’; GFAP: 5’-GGGGCAAAAGCACCAAAGAAG-3’ and 5’-GGGACAACTTGTATTGTGAGCC-3’. The primers used for VCAM, ICAM and P-Selectin were from the QuantiTect primer assay kit (Qiagen). Results are expressed following normalization using 18S RNA.

### Immunofluorescence microscopy analysis

Sampled brains were embedded in OCT (Tissue-Tek, Torrance, CA) and frozen in isopentane at −25°C. 20 μm sections were cut from frozen brains in a cryostat and fixed in 4% paraformaldehyde for 15 min at 21°C. Samples were permeabilzed for 60 min in PBS containing 0.25 % Triton X-100 and 5% newborn goat serum, prior to incubation with primary antibodies at the following dilutions: anti-GFAP (1:500), anti-PN capsular (1:300), and AlexaFLuor 568-IB4 (1:100). Alex-conjugated secondary antibodies and Phalloidin were used at a 1:200 dilution. DAPI was used at a 0.1 mg/ml final concentration. Samples were mounted in DAKO fluorescence mounting medium (DAKO Corp.). fixed samples were analyzed using Eclipse Ti inverted microscopes (Nikon) equipped with a 60 x objective, a CSUX1-A1 spinning disk confocal head (Yokogawa) and a Coolsnap HQ2 camera (Roper Scientific Instruments), or a CSU1-W1 confocal head (Yokogawa) and an ORCA Flash4 CMOS camera (Hamamatsu) controlled by the Metamorph 7.7 software. For live calcein assays and Ca^2+^ imaging, epifluorescence microscopy was performed using a DMRIBe microscope (LEICA microsystems) using 380 nm, 470 nm, or 546 nm LED source excitation, equipped with a Cascade 512 camera (Roper Scientific) driven by the Metamorph (7.7) software. Images were analyzed using the Metamorph software.

### Cell challenge with bacterial strains and pneumolysin

Cultured cells were seeded on sterile 25 mm-diameter coverslips (Deckgläser) at a density of 2 × 10^5^ cells / well in the day before the experiments. Cells were washed 2 times with EM buffer (120mM NaCl, 7mM KCl, 1,8mM CaCl2, 0,8mM MgCl2, 25mM HEPES pH 7.3) supplemented with 5 mM glucose. Cells were incubated with freshly grown bacteria resuspended in EM buffer at a final OD_600nm_ = 0.2, or purified Ply at the indicated concentrations for 90 min at 37°C. Samples were fixed with 3.7% paraformaldehyde, permeabilized by incubation in PBS buffer containing 0.1% Triton X-100 for 4 min at 21°C and processed for immunofluorescence staining of GFAP, F-actin, and bacterial capsule.

### Calcein release assays and Ca^2+^ imaging

Calcein release assays were performed as previously described [30]. Briefly, cultured cells were seeded on sterile 25 mm-diameter coverslips (Deckgläser) at a density of 2 x 10^5^ cells / well in the day before the experiments. Cells were washed 2 times with EM buffer and loaded with calcein-AM at 3 μM final concentration in EM buffer for 30 min at 21°C. Samples were washed three times with EM buffer and placed in a observation chamber on the microscope stage at 37°C. Samples were incubated with purified Ply at 300 nM final concentration and images were acquired at 470 nm excitation every 3 minutes for 60 min. The rates of calcein release were inferred from linear fits with a Pearson correlation coefficient > 0.95.

Ca^2+^ imaging was performed as described previously [31]. Cells were loaded with the fluorescent Ca^2+^ indicator Fluo4-AM at a final concentration of 3 μM for 20 min at 21°C. Cells were washed 3 times with EM buffer and further incubated in EM buffer for 20 min prior to mounting in the observation chamber on the microscope stage. Samples were incubated with Ply at the indicated concentration. Images were acquired every 3 seconds for at least 3 min for each Ply concentration.

### Image analysis

Identical acquisition and grey levels display parameters for all samples from the same set of experiments. A miminimum of 300 fields representing 4.3 x 10^4^ μm^2^ was scored for the determination of bacterial translocation in brain slices following immunofluorescent labeling for PN capsule in at least three independent experiments. The microcolony area was determined from the sum of projected confocal planes subjected to binary thresholding. For analysis of the GFAP filament network in astrocytes from brain slices, GFAP fragments were scored using the “analyze particles “plug-in Fiji using a minimal size threshold value of 0.05 μm^2^ and a circularity index > 0.9 in area proximal to bacterial microcolonies delimited by the presence of detectable shed capsular remnants, or the rest of the sample field. Values are expressed as numbers of GFAP fragments normalized to the surface and are representative of at least 5500 fragments in10 fields from three independent experiments. The area of DAPI-stained nucleus in in vitro grown astrocytes was determined from fluorescent images acquired at a 63x Objective corresponding to a single plane of the epifluorescent DMRIbe microscope, following thresholding.

### Statistical analysis

Statistical difference was analyzed with a non-parametric Wilcoxon test for CFUs determinations, qRT-PCR values, bacterial micocolony area and calcein release assays; Mann-Whitney test for the nuclear area and cell rounding; one-way ANOVA test for the assays involving cell treatment with various Ply concentrations. *: p < 0.05; **: p < 0.01; ***: p < 0.001. ****: p < 0.0001.

## References

1. Mook-Kanamori, B.B., et al., Pathogenesis and pathophysiology of pneumococcal meningitis. Clin Microbiol Rev, 2011. 24(3): p. 557–91.

2. Doran, K.S., et al., Host-pathogen interactions in bacterial meningitis. Acta Neuropathol, 2016. 131(2): p. 185–209.

3. Yau, B., et al., BloodBrain Barrier Pathology and CNS Outcomes in Streptococcus pneumoniae Meningitis. Int J Mol Sci, 2018. 19(11).

4. Iovino, F., et al., How Does Streptococcus pneumoniae Invade the Brain? Trends Microbiol, 2016. 24(4): p. 307–315.

5. Mitchell, T.J. and C.E. Dalziel, The biology of pneumolysin. Subcell Biochem, 2014. 80: p. 145–60.

6. Mook-Kanamori, B., et al., Characterization of a pneumococcal meningitis mouse model. BMC Infect Dis, 2012. 12: p. 71.

7. Iliev, A.I., et al., Cholesterol-dependent actin remodeling via RhoA and Rac1 activation by the Streptococcus pneumoniae toxin pneumolysin. Proc Natl Acad Sci U S A, 2007. 104(8): p. 2897–902.

8. Fortsch, C., et al., Changes in astrocyte shape induced by sublytic concentrations of the cholesterol-dependent cytolysin pneumolysin still require pore-forming capacity. Toxins (Basel), 2011. 3(1): p. 43–62.

9. Wippel, C., et al., Bacterial cytolysin during meningitis disrupts the regulation of glutamate in the brain, leading to synaptic damage. PLoS Pathog, 2013. 9(6): p. e1003380.

10. Grab, D.J., et al., How can microbial interactions with the blood-brain barrier modulate astroglial and neuronal function? Cell Microbiol, 2011. 13(10): p. 1470–8.

11. Abbott, N.J., et al., Structure and function of the blood-brain barrier. Neurobiol Dis, 2010. 37(1): p. 13–25.

12. Abbott, N.J., L. Ronnback, and E. Hansson, Astrocyte-endothelial interactions at the blood-brain barrier. Nat Rev Neurosci, 2006. 7(1): p. 41–53.

13. Iadecola, C., The Neurovascular Unit Coming of Age: A Journey through Neurovascular Coupling in Health and Disease. Neuron, 2017. 96(1): p. 17–42.

14. Aspelund, A., et al., A dural lymphatic vascular system that drains brain interstitial fluid and macromolecules. J Exp Med, 2015. 212(7): p. 991–9.

15. Boulay, A.C., S. Cisternino, and M. Cohen-Salmon, Immunoregulation at the gliovascular unit in the healthy brain: A focus on Connexin 43. Brain Behav Immun, 2016. 56: p. 1–9.

16. Yardeni, T., et al., Retro-orbital injections in mice. Lab Anim (NY), 2011. 40(5): p. 155–60.

17. Kietzman, C.C., et al., Dynamic capsule restructuring by the main pneumococcal autolysin LytA in response to the epithelium. Nat Commun, 2016. 7: p. 10859.

18. Braun, J.S., et al., Pneumolysin causes neuronal cell death through mitochondrial damage. Infect Immun, 2007. 75(9): p. 4245–54.

19. Giaume, C., et al., Connexin and pannexin hemichannels in brain glial cells: properties, pharmacology, and roles. Front Pharmacol, 2013. 4: p. 88.

20. De Bock, M., L. Leybaert, and C. Giaume, Connexin Channels at the Glio-Vascular Interface: Gatekeepers of the Brain. Neurochem Res, 2017. 42(9): p. 2519–2536.

21. Stringaris, A.K., et al., Neurotoxicity of pneumolysin, a major pneumococcal virulence factor, involves calcium influx and depends on activation of p38 mitogen-activated protein kinase. Neurobiol Dis, 2002. 11(3): p. 355–68.

22. Skals, M., et al., Alpha-hemolysin from Escherichia coli uses endogenous amplification through P2X receptor activation to induce hemolysis. Proc Natl Acad Sci U S A, 2009. 106(10): p. 4030–5.

23. Skals, M., et al., Bacterial RTX toxins allow acute ATP release from human erythrocytes directly through the toxin pore. J Biol Chem, 2014. 289(27): p. 19098–109.

24. Tran Van Nhieu, G., et al., Connexin-dependent inter-cellular communication increases invasion and dissemination of Shigella in epithelial cells. Nat Cell Biol, 2003. 5(8): p. 720–6.

25. van Pee, K., et al., CryoEM structures of membrane pore and prepore complex reveal cytolytic mechanism of Pneumolysin. Elife, 2017. 6.

26. El-Rachkidy, R.G., N.W. Davies, and P.W. Andrew, Pneumolysin generates multiple conductance pores in the membrane of nucleated cells. Biochem Biophys Res Commun, 2008. 368(3): p. 786–92.

27. Gilbert, R.J. and A.F. Sonnen, Measuring kinetic drivers of pneumolysin pore structure. Eur Biophys J, 2016. 45(4): p. 365–76.

28. Wang, Y. and S. Wang, Increased extracellular ATP: an omen of bacterial RTX toxin-induced hemolysis? Toxins (Basel), 2014. 6(8): p. 2432–4.

29. Boulay, A.C., et al., Immune quiescence of the brain is set by astroglial connexin 43. J Neurosci, 2015. 35(10): p. 4427–39.

30. Guignot, J., A. Segura, and G. Tran Van Nhieu, The Serine Protease EspC from Enteropathogenic Escherichia coli Regulates Pore Formation and Cytotoxicity Mediated by the Type III Secretion System. PLoS Pathog, 2015. 11(7): p. e1005013.

31. Tran Van Nhieu, G., et al., Actin-based confinement of calcium responses during Shigella invasion. Nat Commun, 2013. 4: p. 1567.

