## supplemental Figs 1-7 for "Role of astrogial Connexin 43 in pneumococcal meningitis and pneumolysin cytotoxicity"

**Supplementary information**

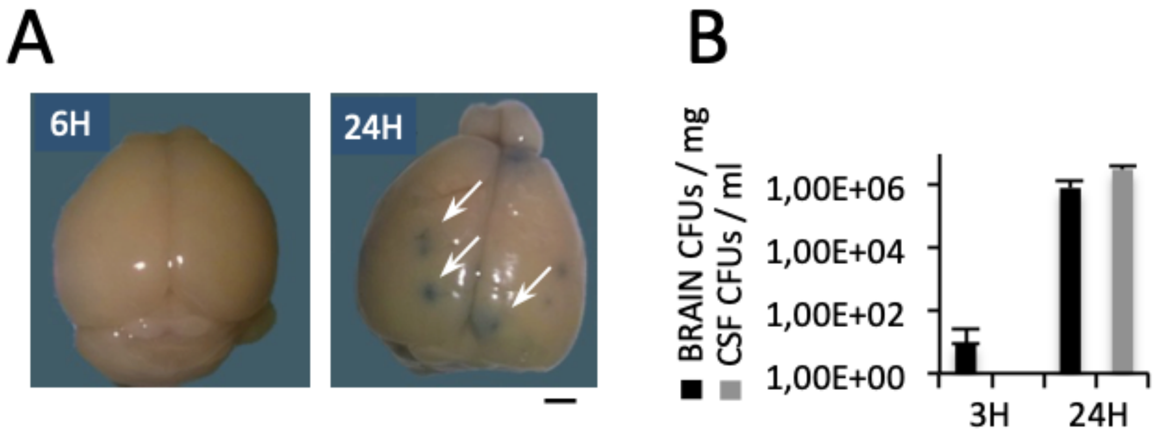


**Fig. S1. PN meningitis in mice following retro-orbital vein injection.**

6-9 weeks old C57BL/6 mice were infected through intravenous retro-orbital with $10^7$ bacterial CFUs. **A**, at the indicated times, mice were subjected wih intracardiac perfusion with buffer than Blue Evans-containing buffer prior to brain sampling. The arrows indicate Blue Evans leakage associated with macroscopic sites of BBB rupture. Scale bar = 1 mm. **B**, bacterial CFU determination in: B, brain (solid bars); cerebrospinal fluid (grey bars). N = 3, > 3 mice per determination.

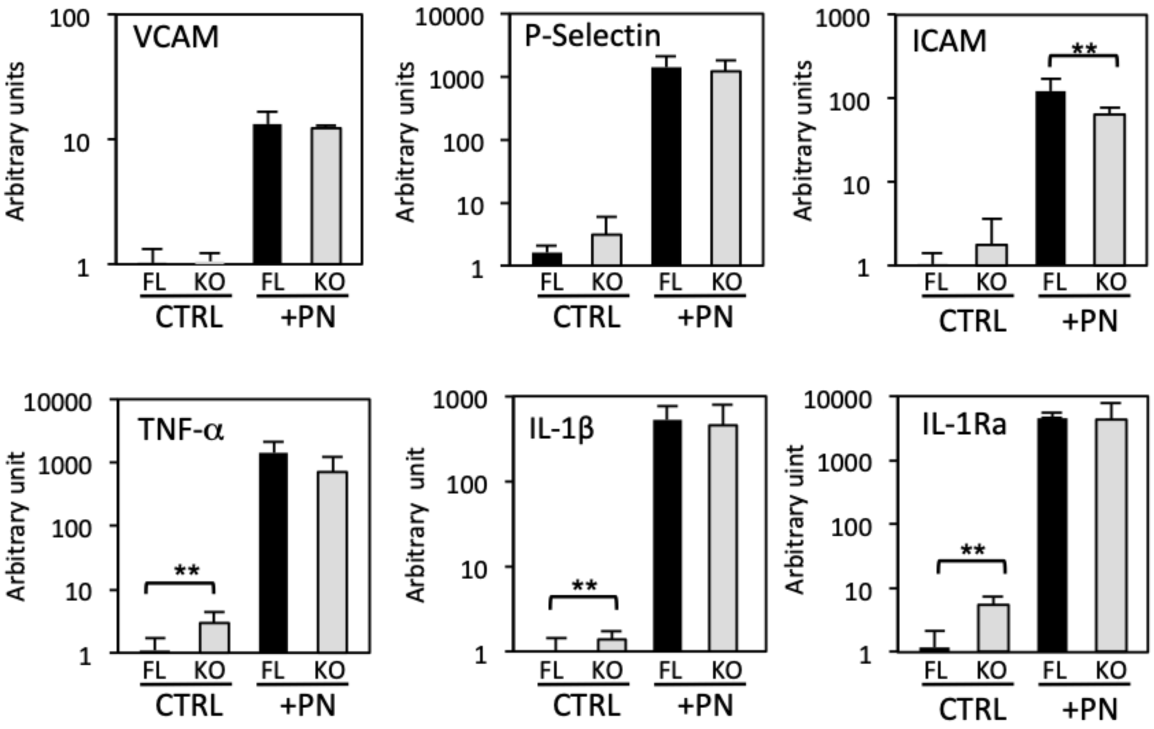


**Fig. S2. Up-regulation of inflammatory markers during PN infection.**

6-9 weeks old C57BL/6 mice were infected through intravenous retro-orbital with $10^7$ bacterial CFUs. At 13H post-infection, qRT-PCR was performed on total RNAs extracted from brain samples using primers specific to the indicated markers (Methods). Results are expressed as average determination value in arbitrary units normalized to values obtained for 16S mRNA. CTRL: uninfected mice. + PN: mice challenged with PN. FL: mice expressing aCx43; KO: aCx43$^{-/-}$ mice. FL: N = 6, 6 mice per determination; KO: N = 6, 6 mice per determination. Mann-Whitney. **: $p < 0.01$.

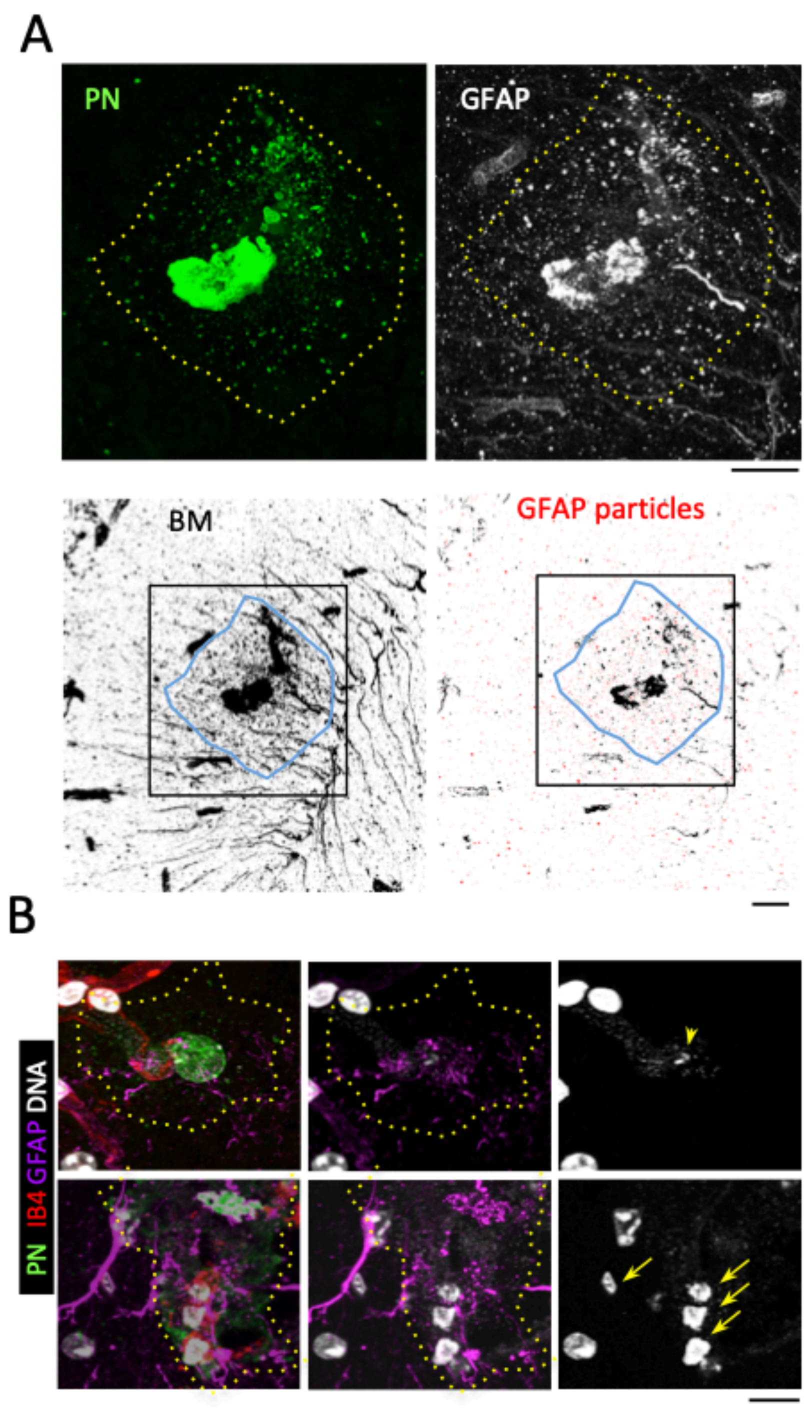


**Fig. S3. GFAP fragmentation and nuclear shrinkage during PN meningitis.**

6-9 weeks old C57BL/6 mice were infected through intravenous retro-orbital with $10^7$ bacterial CFUs. Brains were sampled at 13H post-infection and 20 um section brain slices were processed for immunofluorescence staining. Scale bar = 5 µm. **A**, top panels: sum of projected confocal planes corresponding to a higher magnification of the boxed insets shown in lower panels. Green: PN capsule; gray levels: GFAP. Yellow dotted outline: area associated with the PN microcolony showing capsular remnants. Bottom panels, blue outline:

area associated with the PN microcolony showing capsular remnants shown in yellow in top panels. BM: binary mask of GFAP staining. GFAP particles: detected GFAP particles are outlined in red (Methods). **B**, red: IB-4 endothelial staining; green: PN capsule; gray levels: DNA. Yellow dotted outline: area associated with the PN microcolony showing capsular remnants. Arrowhead: nuclear fragmentation. Arrows: nuclear shrinkage.

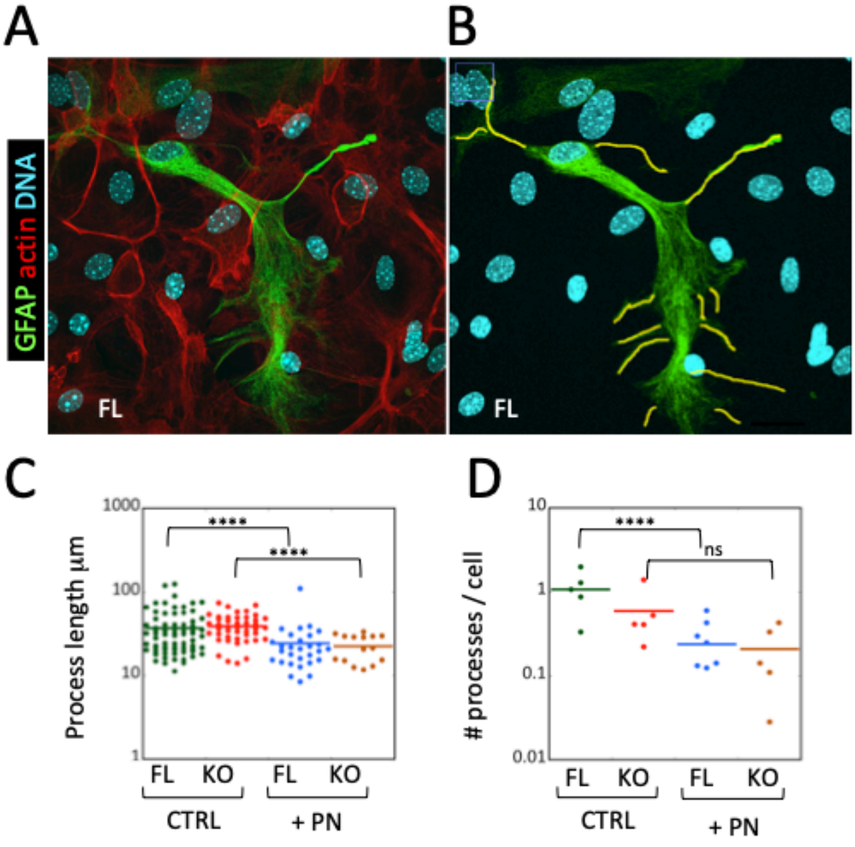


**Fig. S4. Quantification of astrocytic processes during PN meningitis.**

*In vitro* cultured astrocytes derived from the brain cortexes of mice expressing aCx43 (FL) or aCx43$^{-/-}$ mice (KO) were challenged with wild-type TIGR4 (PN) and processed for immunofluorescence staining. **A**, **B**, staining for: red: F-actin; green: GFAP; cyan: DNA. **B**, astrocytic processes outlined in yellow. Scale bar = 5 µm. C, D, CTRL: uninfected cells. + PN: cells challenged with PN. FL: cells expressing aCx43; KO: aCx43$^{-/-}$ cells. **C**, median length of astrocytic processes. **D**, median number of GFAP processes per cell. FL: N = 2, 78 cells; KO: N = 2, 93 cells. FL+PN: N = 2, 131 cells; KO+PN: N = 2, 103 cells. Mann-Whitney. ****: $p < 0.0001$.

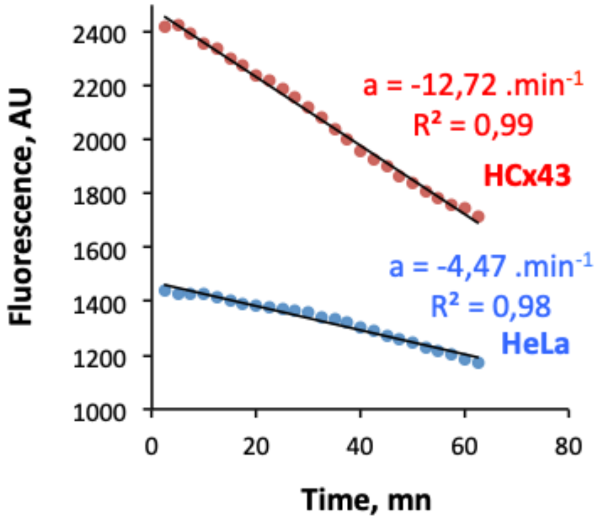


**Fig. S5. Role of Cx43 in Ply-mediated calcein release.**

Calcein loaded cells were challenged with Ply at 250 nM final concentration. Traces are representative of the variation of average calcein fluorescence intensity in single cells for HeLa (blue) and HCx43 (red) cells in arbitrary units (AU). Solid lanes: linear fits. The rate of fluorescence intensity decrease (a) and Pearson correlation coefficient ($R^2$) are shown with the corresponding color code.

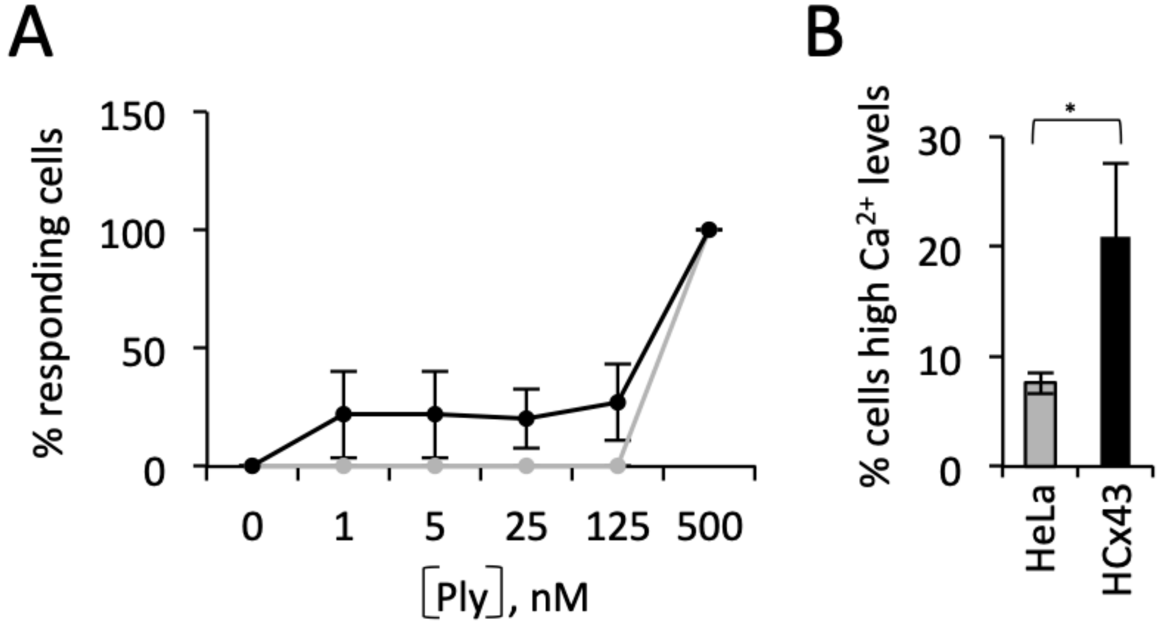


**Fig. S6. $Ca^{2+}$ responses to Ply in HeLa and HCx43 cells**

Cells were loaded with Fluo-4 and challenged with Ply at the indicated concentrations. **A**, percent of cells showing $Ca^{2+}$ responses at the indicated Ply concentration. Grey circles: HeLa cells; N = 3, 145 cells. Solid circles: HCx43 cells. N = 107 cells. **B**, percent of cells with lasting high Ca2+ levels following treatment with 500 nM Ply. Grey bar: HeLa cells, N = 3, 130 cells. Solid bar: HCx43 cells, N = 3, 89 cells. Kruskal-Wallis. *: p = 0.049.

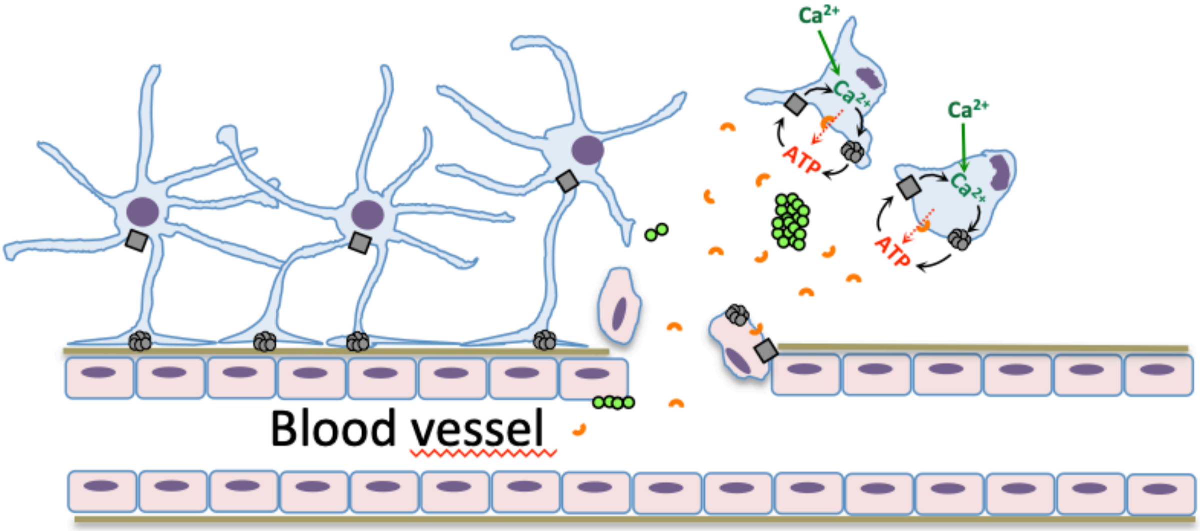


**Fig. S7. Role of aCx43 during PN meningitis.**

Astrocytes regulate the BBB function through their end-feet contacting blood vessels. Crossing of the BBB by PN is associated with the destabilization of blood vessels seals. Secreted Ply induces the release of ATP (red arrow) in the extracellular space by forming small pores or “arcs” in the plasma membrane of astrocytes (orange half circles). Black arrows: ATP stimulates $Ca^{2+}$ signaling via purinergic receptors (grey box). Increase cytosolic $Ca^{2+}$ activates the opening of Cx43 hemichannels (grey circles) further amplifying ATP release and cytosolic $Ca^{2+}$ increase, resulting in the destruction of astrocytic processes, plasma membrane permeabilization and eventually astrocytic death linked to $Ca^{2+}$ influx (green arrow) and overload. Disruption of astrocyte-endothelial cells regulation fragilizes the BBB, favoring PN translocation and growth in the brain cortex due to local perfusion of blood vessel lumenal content.

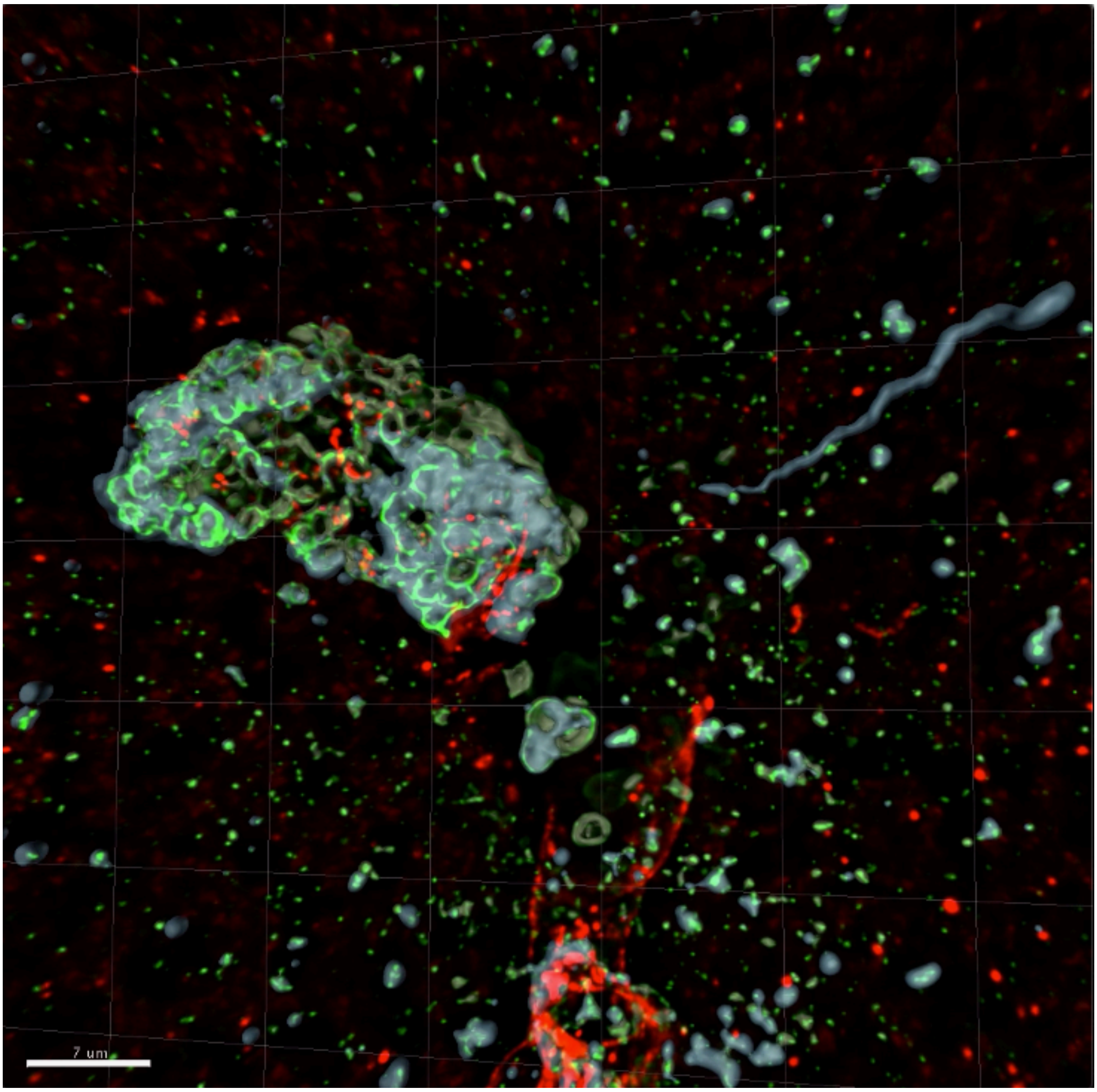


**Movie. S1. 3D-Surface rendering of GFAP fragmentation during PN meningitis.** 3D-reconstruction was performed following deconvolution of confocal planes using the Huygens software. Surface rendering on was performed using the Imaris software. Green: PN capsule; grey: GFAP; red: IB-4. Scale bar = 7 µm.
